# Segregational instability of multicopy plasmids: a population genetics approach

**DOI:** 10.1101/2022.03.15.484385

**Authors:** V Miró Pina, JCR Hernández, A Siri-Jégousse, R Peña-Miller, S Palau, A González Casanova

## Abstract

Plasmids are extra-chromosomal genetic elements that encode a wide variety of phenotypes and can be maintained in bacterial populations through vertical and horizontal transmission, thus increasing bacterial adaptation to hostile environmental conditions like those imposed by antimicrobial substances. To circumvent the segregational instability resulting from randomly distributing plasmids between daughter cells upon division, non-transmissible plasmids tend to be carried in multiple copies per cell, with the added benefit of exhibiting increased gene dosage and resistance levels. But carrying multiple copies also results in a high metabolic burden to the bacterial host, therefore reducing the overall fitness of the population. This trade-off poses an existential question for plasmids: What is the optimal plasmid copy number? In this manuscript, we address this question by postulating and analyzing a population genetics model to evaluate the interaction between selective pressure, the number of plasmid copies carried by each cell, and the metabolic burden associated with plasmid bearing in the absence of selection for plasmid-encoded traits. Parameter values of the model were estimated experimentally using *Escherichia coli* K12 carrying a multicopy plasmid encoding for a fluorescent protein and *bla*_TEM-1_, a gene conferring resistance to *β*-lactam antibiotics. By numerically determining the optimal plasmid copy number for constant and fluctuating selection regimes, we show that plasmid copy number is a highly optimized evolutionary trait that depends on the rate of environmental fluctuation and balances the benefit between increased stability in the absence of selection with the burden associated with carrying multiple copies of the plasmid.

## 1 Introduction

Prokaryotes transfer DNA at high rates within microbial communities through mobile genetic elements such as bacteriophages,^1^ transposons^2^ or extra-chromosomal DNA molecules known as plasmids.^3^ Crucially, plasmids have core genes that allow them to replicate independently of the chromosome but also encode for accessory genes that provide their bacterial hosts with new functions and increased fitness in novel or stressful environmental conditions.^4^ Plasmids have been widely studied due to their biotechnological potential^5^ and their relevance in agricultural processes,^6^ but also because of their importance in clinical practice since they have been identified as significant factors contributing to the current global health crisis generated by drug resistant bacterial pathogens.^7^

Although the distribution of plasmid fitness effects is variable and context dependant,^8^ it is generally assumed that in the absence of selection for plasmid-encoded genes, plasmids impose a fitness burden on their bacterial hosts.^9, 10^ As a result, plasmid-bearing populations can have a competitive disadvantage compared to plasmid-free cells, thus threatening plasmids to be cleared from the population through purifying selection.^11^ To avoid extinction, some plasmids can transfer horizontally to lineages with increased fitness, with previous theoretical results establishing sufficient conditions for plasmid maintenance, namely that the rate of horizontal transmission has to be larger than the combined effect of segregational loss and fitness cost.^12, 13^ Also, some plasmids encode molecular mechanisms that increase their stability in the population, for instance, toxin-antitoxin systems that kill plasmid-free cells,^14^ or active partitioning mechanisms that ensure the symmetric segregation of plasmids upon division.^15^

To avoid segregational loss, non-conjugative plasmids lacking active partitioning and post-segregational killing mechanisms tend to be present in many copies per cell, therefore decreasing the probability of producing a plasmid-free cell when randomly segregating plasmids during cell division. But this reduced rate of segregational loss is not sufficient to explain the stable persistence of costly plasmids in the population, suggesting that a necessary condition for plasmids to persist in the population is to carry beneficial genes for their hosts that are selected for in the current environment. However, regimes that positively select for plasmid-encoded genes can be sporadic and highly specific, so plasmid persistence is not guaranteed in the long term. Moreover, even if a plasmid carries useful genes for the host, these can be captured by the chromosome, thus making plasmids redundant and rendering them susceptible to be cleared from the population.^16^ This evolutionary dilemma has been termed the ‘plasmid paradox’.^17^

In this paper, we use a population genetics modeling approach to evaluate the interaction between the number of plasmid copies contained in each cell and the energetic cost associated with carrying each plasmid copy. We consider a non-transmissible, multicopy plasmid (it can only be transmitted vertically) that lacks active partitioning or post-segregational killing mechanisms (plasmids segregate randomly upon division). We will also consider that plasmids encode a gene that increases the probability of survival to an otherwise lethal concentration of an antimicrobial substance, albeit imposing a burden to plasmid-bearing cells in drug-free environments. To estimate parameters of our population genetics model, we used an experimental model system consisting on *Escherichia coli* bearing a multicopy plasmid pBGT (∼19 copies per cell) carrying *bla*_TEM-1_, a drug-resistance gene that produces a *β* -lactamase that degrades ampicillin and other *β* -lactam antibiotics.^7, 18^

We used computer simulations to evaluate the stability of a multicopy plasmid in terms of the duration and strength of selection in favor of plasmid-encoded genes. This allowed us to numerically estimate the number of copies that maximized plasmid stability under different environmental regimes: drug-free environments, constant exposure to a lethal drug concentration, and intermittent periods of selection. Altogether, our results confirm the existence of two opposing evolutionary forces acting on the number of copies carried by each cell: selection against high-copy plasmids consequence of the fitness cost associated with bearing multiple copies of a costly plasmid and purifying selection resulting from the increased probability of plasmid loss observed in low-copy plasmids.

## Methods

### 1.1 Serial dilution protocol

We consider a serial dilution experiment with two types of bacteria: plasmid-bearing (PB) and plasmid-free (PF). Let us denote by *n* the plasmid copy number (PCN) and argue that this is an important parameter: in the one hand, the selective disadvantage of PB individuals due to the cost of carrying plasmids is assumed to be proportional to *n*; on the other hand, the PCN determines the heritability of the plasmid.

In our schema, each day starts with a population of *N* cells that grow exponentially until saturation is reached (i.e. until there are *γN* cells). At the beginning of the next day, *N* cells are sampled (at random), transferred to new media and exponential growth starts again (Figure 1A).

**Figure 1.**
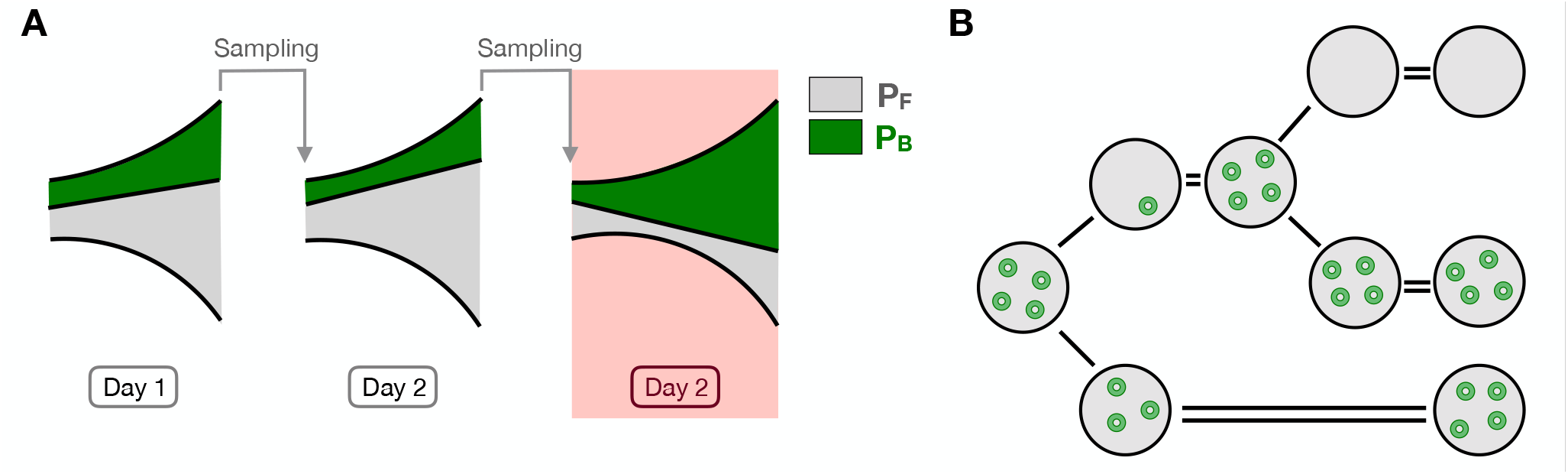
Schematic diagram of the model. **A)** Serial dilution protocol. PB cell are represented in green while PF cells are represented in gray. We show three days of the experiments. An antibiotic pulse is added during day 3. **B)** Segregational loss. Upon cell division, plasmids are segregated at random between the two daughter cells. Then the plasmids are replicated until the PCN is 4. When a cell inherits no plasmid, it becomes plasmid free.

### 1.2 Inter-day dynamics

To model the inter-day dynamics, we consider a discrete-time model in which the population size is fixed to *N*. Day *i* starts with a fraction *X*_*i*_ of PB cells (and 1 − *X*_*i*_ of PF cells). We consider that the fitness cost associated with plasmid maintenance, *κ*_*n*_ is proportional to the PCN, i.e. *κ*_*n*_ = *κn*. This means that, at the end of day *i*, the number of PF cells is proportional to their initial frequency 1 − *X*_*i*_, while the number of PB cells is proportional to their initial frequency *X*_*i*_ multiplied by (1 − *κ*_*n*_) < 1. So, at the end of day *i*, the fraction of PB cells would be

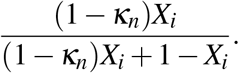

In addition, PB cells can lose their plasmids an become PF and with probability *μ*_*n*_, so, at the end of day *i*, the fraction of PB cells needs to be multiplied by (1 − *μ*_*n*_).

At the beginning of day *i* + 1, we sample *N* individuals at random from the previous generation. Since *N* is very large, we can neglect stochasticity and assume that the fraction of PB cells at the beginning of day *i* + 1 is equal to their fraction at the end of day *i*, i.e.

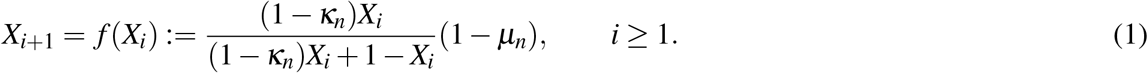

Additionally, we aim to modeling selection for plasmid-encoded genes. For plasmids carrying antibiotic resistance genes, this is achieved by exposing the population to antibiotic pulses. Individuals with no plasmids suffer more from this this treatment so, at each pulse, we observe an increment in the relative frequency of the PB subpopulation. To model this phenomenon, we assume that, in the presence of antibiotic, PF individuals exhibit a selective disadvantage represented by parameter *α* ∈ [0, 1].

For instance, if an antibiotic pulse occurs at day *i*, all PB cells survive, (there are *NX*_*i*_), but the PF cells die with probability *α*, so only *N*(1 − *α*)(1 − *X*_*i*_) survive. So, the fraction of PB individuals, right after the antibiotic pulse becomes

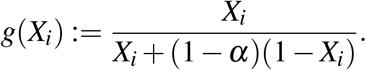

Then, cells grow exponentially again, as in a normal day, so that, at the end of the day, the fraction of PB cells is *f* (*g*(*X*_*i*_)).

If we consider that the pulses occur at generations *T*, 2*T*,…, the frequency process becomes

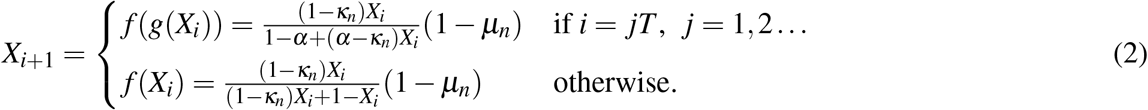

### 1.3 Intra-day dynamics

For the intra-day dynamics, day *i* starts with a population of *N* cells (*N* ∼ 10^5^ in the experiment) that grow exponentially until saturation is reached (i.e. until there are *γN* cells.). The initial fraction of PB cells is *X*_*i*_. We assume that, in the absence of antibiotic, the population evolves as a continuous time multi-type branching process 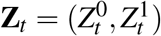, where 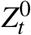 (resp 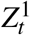) is the number of PB cells (resp. PF cells). The reproduction rate (or *Malthusian fitness*) of PB (resp. PF) individuals is *r* (resp. *r* + *ρ*_*n*_), with *ρ*_*n*_ > 0 (since PB individuals have some disadvantage due to the cost of plasmid maintenance). Following^19^, we assume that *ρ*_*n*_ ∼ *N*^−*b*^ for some *b* ∈ (0, 1/2) (this regime is known as *moderate-strong selection*).

We consider plasmids that lack active partitioning systems^20^ so, at the moment of cell division, each plasmid randomly segregates into one of the two new cells. Once in the new host, the plasmids replicates until reaching *n* copies. If, however, one of the two new cells has all the *n* copies, the other one will not carry any plasmid copy and becomes PF. Thus, we make the simplifying assumption that the daughter of a PB cell becomes PF with probability 2^−*n*^ (segregational loss rate), as illustrated in Figure 1B. Therefore, at every branching event, an individual splits in two. Plasmid-free individuals only split in two PF individuals. Plasmid-bearing individuals can split in one PF individual and one PB individual with probability 2^−*n*^ (if all the plasmids go to one of them) or they can split in two plasmid-bearing individuals with probability 1 − 2^−*n*^.

Let *M*(*t*) = {*M*_*i, j*_(*t*) : *i, j* = 0, 1} be the mean matrix given by 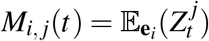, the average size of the type *j* population at time *t* if we start with a type *i* individual. According to [21, Section V.7.2], *M*(*t*) can be calculated as an exponential matrix

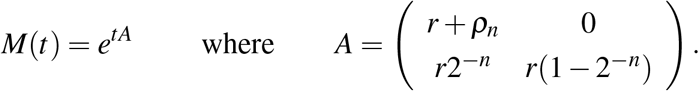

More precisely,

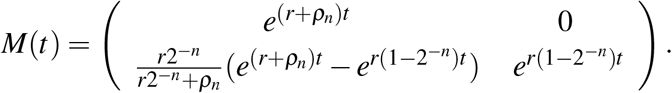

Let *σ* be the duration of the growth phase. Since *N* is very large, one can assume that reproduction is stopped when the expectation of the number of descendants reaches *γN*, i.e. that *σ* satisfies

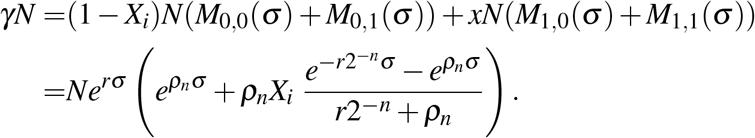

Since *ρ*_*n*_ ∼ *N*^−*b*^, we have for large enough *N* that

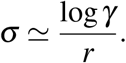

Since γ*N* >> 1, we can assume that the number of PB (resp. PF) cells at the end of the day is equal to its expected value. Therefore, the fraction of PB cells at the end of day *i* is equal to

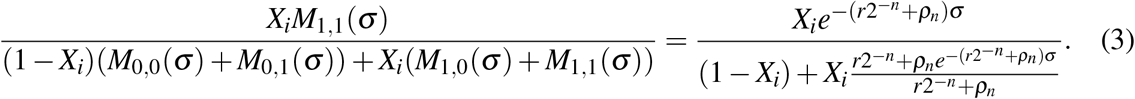

This corresponds to equation (2) with parameters

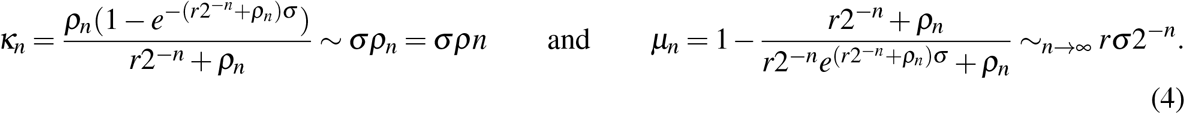

The importance of these formulas is that they connect measurable quantities with theoretical parameters, leading to a method to estimate the parameters of the model from experiments, which is the spirit of the experiment described in the following section.

### 1.4 Model parametrization

Our goal is to use the inter-day model to evaluate the long-term dynamics of plasmid-bearing populations in terms of the cost associated with carrying plasmids and the fitness advantage conferred by the plasmid in the presence of positive selection. To quantify these parameters experimentally, our approach consisted in two phases: (1) from growth kinetic experiments, we estimate parameters *ρ, r* and *σ* of the inter-day model, and (2) we perform competition experiments in a range of drug concentrations to obtain *μ*_*n*_ and *κ*_*n*_ using equation (4) of the intra-day model.

Our experimental model system consisted in *Escherichia coli* K12 carrying pBGT, a non-transmissible multicopy plasmid used previously to study plasmid dynamics and drug resistance evolution.^22–25^ Briefly, pBGT is a ColE1-like plasmid with ∼19 plasmid copies per cell, lacking the necessary machinery to perform conjugation or to ensure symmetric segregation of plasmids upon division. This plasmid carries a GFP reporter under an arabinose-inducible promoter and the *bla*_TEM-1_ gene that encodes for a *β* -lactamase that efficiently degrades *β* -lactam antibiotics, particularly ampicillin (AMP). The minimum inhibitory concentration (MIC) of PB cells to AMP is 8, 192 mg/l, while the PF strain has a MIC of 4 mg/l (see Appendix A).

Growth experiments were performed in 96-well plates with lysogeny broth (LB) rich media and under controlled environmental conditions. Using an plate absorbance spectrophotometer, we obtained bacterial growth curves that enabled us to estimate the maximal growth rate of the PB and PF strains, corresponding to *r* and *ρ*_*n*_ in the intra-day model^26^ (Figure 2A and Appendix C). As expected, we observed a reduction in bacterial fitness of the PB subpopulation, expressed in terms of a decrease in its maximum growth rate when grown in isolation. The metabolic burden associated with carrying the pBGT plasmid (*n* = 19) was estimated at 0.108 ± 0.067 (Figure 2B).

**Figure 2.**
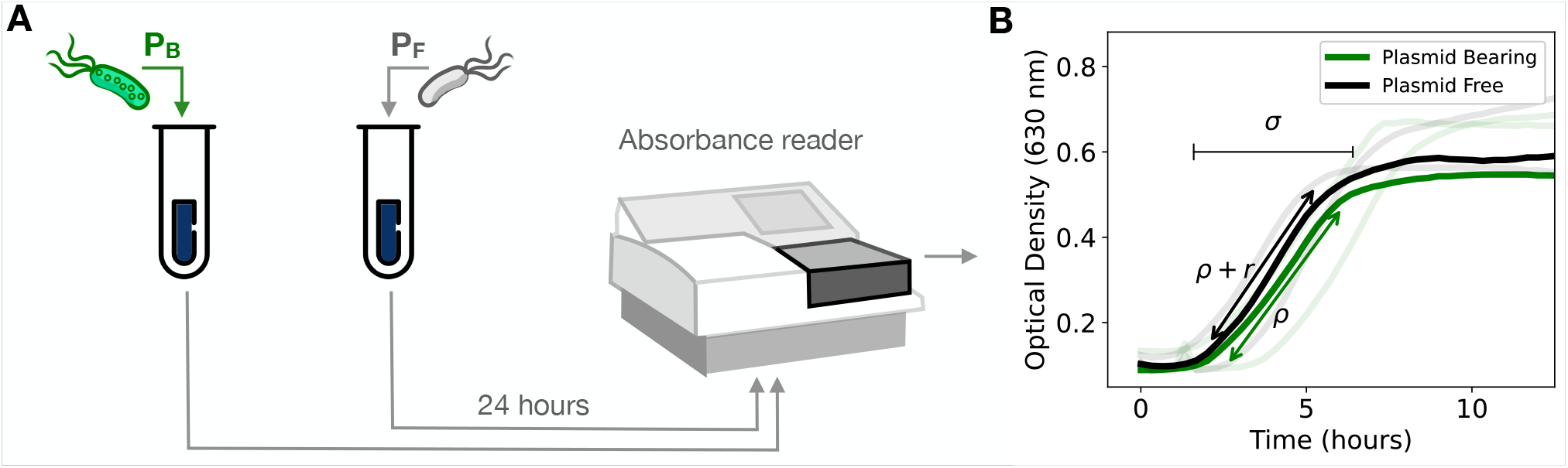
Growth kinetic experiment. **A)** Schematic diagram illustrating a bacterial growth experiment performed in drug-free media separately for PB and PF populations. We used an absorbance microplate reader to measure the optical density (OD_630_) at different time-points during the 24-hour experiment. **B)** Growth curves of PB (green) and PF (black) strains, with replicate experiments represented as shaded curves. The duration of the exponential phase, *σ*, was estimated by identifying the start of exponential phase and the time elapsed before reaching carrying capacity. Parameter *ρ* refers to the maximum growth rate of the PB population, while the selective advantage of the PF strain is represented with *r*.

We then performed a one-day competition experiment consisting of mixing PB and PF subpopulations with a range of relative abundances and exposing the mixed populations to environments with increasing drug concentrations (see Figure 3A for a schematic of the experimental protocol). Previous studies have used a similar approach to determine a selection coefficient,^27^ a quantity that was used to show that selection of resistance can occur even at sub-lethal antibiotic concentrations.^28^ Figure 3B shows the final PF frequency obtained for different initial population structures and strengths of selection.

**Figure 3.**
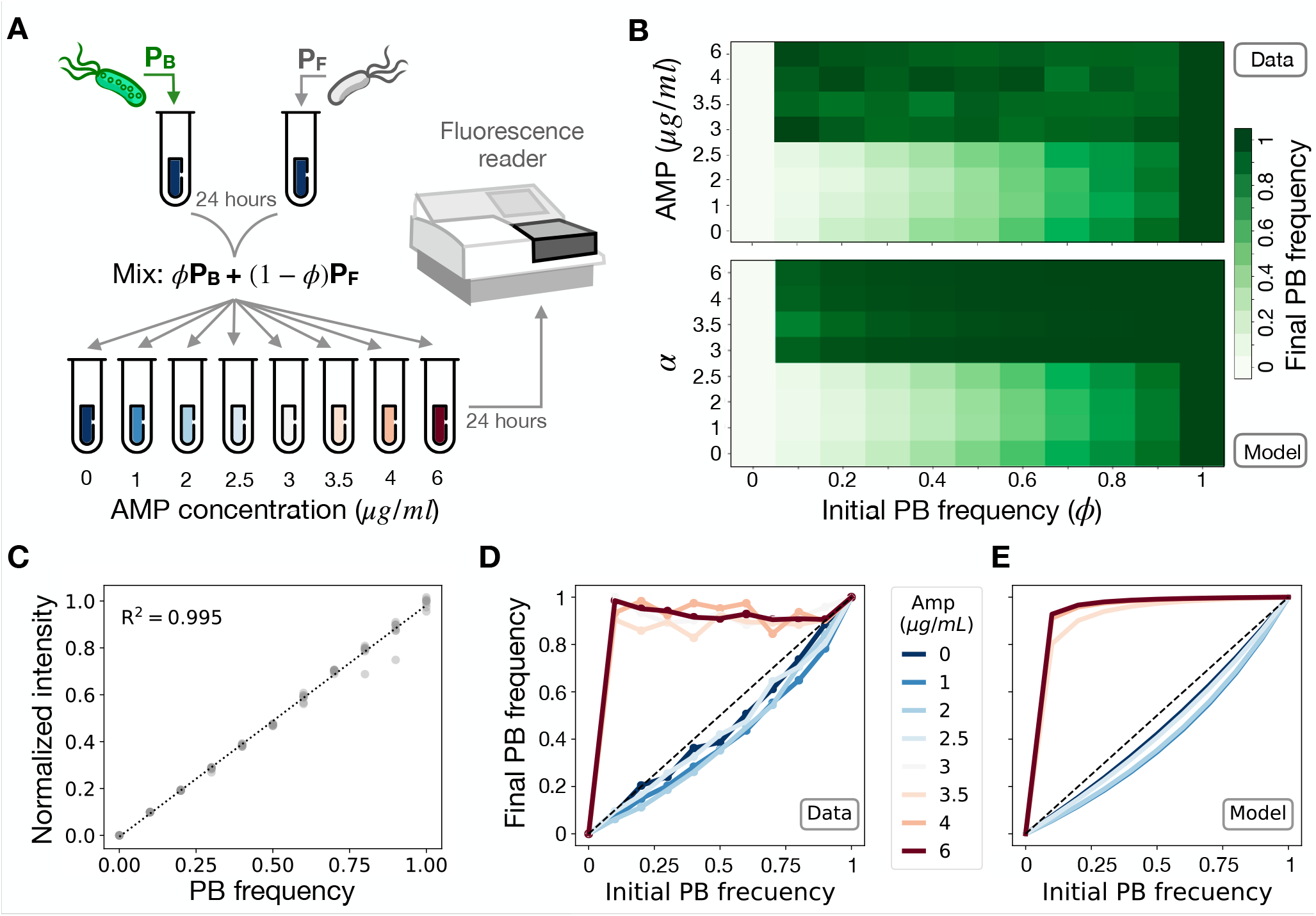
Competition experiment under a range of drug concentrations. **A)** Schematic diagram illustrating an experiment where PB and PF are mixed at different relative abundances and submitted to a range of ampicillin concentrations (0, 1, 2, 2.5, 3, 3.5, 4, and 6 *μ*g/ml). We use a fluorescence spectrophotometer to estimate the relative abundance of plasmid bearing cells in the population after 24 hours of growth. **B)** Final PF frequency (illustrated in a gradient of green) for different initial fraction of PB cells and selection coefficients (top: data; bottom: model). **C)** Control experiment illustrating that normalized fluorescence intensity is correlated with the fraction of the population carrying plasmids. Each dot present a replica and the dotted line a linear regression (R^2^=0.995). **D)** Experimental iterative map showing the existence of a minimum drug concentration that rescues the PB population (red lines). At low drug concentrations (blue lines), the PB population decreases in frequency. **E)** Theoretical iterative map obtained by numerically solving equation (2) for a range of strength of selections and initial PB frequencies. By fixing *κ* _*n*_ (previously estimated by growing each strain in monoculture), we fitted parameter a in equation (2) to the experimental data. Colors indicate the strength of selection (in blue, values of α where the cost of carrying plasmids is stronger than the benefit resulting from positive selection, yielding curves below the identity line. Red curves represent simulations obtained with values of α strong enough to kill PF cells, thus increasing PB frequency in the population.

The fitness cost associated with carrying plasmids in our inter-day model was estimated from the proportion of PB cells at the end of a competition experiment. This quantity can be obtained from the normalized fluorescent intensity of the bacterial culture, measured with a fluorescent spectrophotometer or with flow cytometry (Figure 3C shows a linear relationship between both quantities). Figure 3D shows the end-point bacterial density resulting from competition experiments with different initial fractions of PB cells exposed to a range of AMP concentrations. Note that, at low AMP concentrations (blue lines), the frequency of plasmid-bearing is below the identity, consistent with plasmids imposing a fitness cost to PB cells. In contrast, at high AMP concentrations (red lines), plasmid-free cells are killed and the population is almost exclusively conformed by PB cells.

In the model, since PCN is a fixed parameter, the PB fraction resulting from a competition experiment in the absence of selection only depends on the cost associated with plasmid bearing. Therefore, by fitting equation (1), we estimated that the cost associated with carrying *n* = 19 copies of pBGT was *κ*_*n*_ = 0.272. Furthermore, by fixing this parameter and incorporating antibiotics, we estimated the selective pressure *α* for different antibiotic concentrations by fitting equation (2) to the experimental data. Figure 3E illustrates that at low antibiotic concentrations (small values of *α*) the frequency of the population is low, while higher values of *α* result in an increased PB frequency. Table 1 summarizes parameter values estimated for each strain in our model, and Table 2 shows the correspondence between antibiotic concentrations and *α*.

**Table 1.**
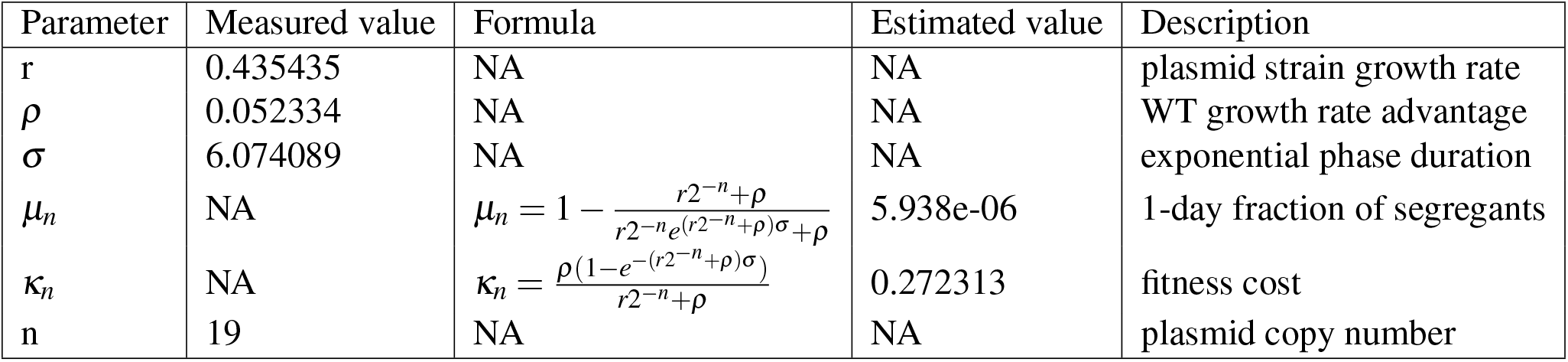
Model parameters estimated using growth curves experiments in the absence of antibiotics.

**Table 2.** Model parameter estimated by fitting equation (4) to experimental data obtained for a range of ampicillin concentrations.

| Amp | $\kappa_n$ | $\alpha$ |
| --- | --- | --- |
| 0.0 | 0.272276 | 0.0 |
| 1.0 | 0.272276 | -0.37781 |
| 2.0 | 0.272276 | -0.332662 |
| 2.5 | 0.272276 | -0.058457 |
| 3.0 | 0.272276 | 0.992911 |
| 3.5 | 0.272276 | 0.9801 |
| 4.0 | 0.272276 | 0.992075 |
| 6.0 | 0.272276 | 0.99373 |

## 2 Results

### 2.1 Segregational instability in the absence of selection

Our first aim was to evaluate the stability of a costly multicopy plasmid in the absence of selection for plasmid-encoded genes (i.e. without antibiotics). By numerically solving equation (1), we evaluated the stability of the PB subpopulation in terms of the mean PCN and the fitness cost associated with carrying each plasmid copy (see Appendix C). As expected, in the absence of selection, plasmids are always cleared from the population with a decay rate that depends on PCN. We define the time-to-extinction as the time when the fraction of PB cells goes below an arbitrary threshold.

For cost-free plasmids (i.e. when *κ* = 0), the time-to-extinction appears to be correlated to PCN (Figure 4A). In contrast, if we consider a costly plasmid (*κ >* 0) and that the total fitness cost is proportional to the PCN (i.e. if *PCN* = *n*, the total cost is *κ*_*n*_ = *κn*), then extinction occurs in a much faster timescale (Figure 4B – notice the difference of timescales with Figure 4A). As shown in Figure 4B, small PCN values are associated with a high probability of segregational loss, and therefore the time-to-extinction increases with PCN. However, large values of PCN are associated with higher levels of instability due to the detrimental effect on host fitness resulting from carrying multiple copies of a costly plasmid.

**Figure 4.**
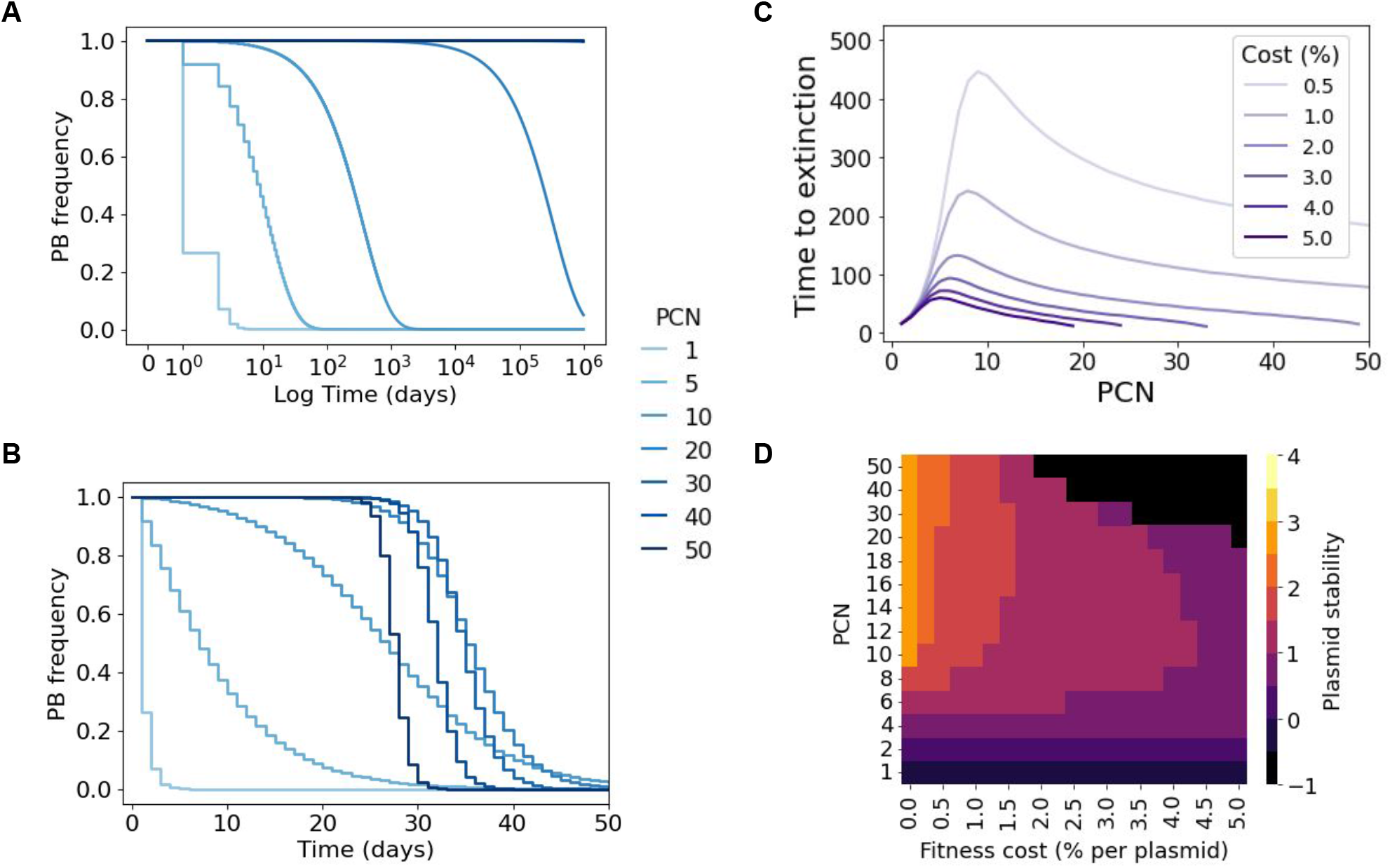
Numerical results for the model without selection for plasmid-encoded genes. **A)** Plasmid frequency as a function of time for a cost-free plasmid (*κ* = 0). Note how, as the PCN increases, the stability of plasmids also increases, although eventually all plasmids will be cleared from the system. **B)** Dynamics of plasmid loss for strains bearing a costly plasmid (*κ* = 0.0143). In this case, low-copy plasmids (light blue lines) are highly unstable, but so are high-copy plasmids (dark blue lines). **C)** Time elapsed before plasmid extinction for a range of PCNs. A very costly plasmid (*κ* = 5%) is represented in dark purple, while the light purple line denotes a less costly plasmid (*κ* = 0.5%). **D)** Plasmid stability for a range of fitness costs and PCNs (discrete colormap indicates level of stability, yellow denotes higher stability, while dark purple denotes rapid extinction). Stability is measured as the area under the curve (AUC) of trajectories similar to those in **B**, expressed in log_10_ scale. Notice that, for intermediate fitness costs, the PCN that maximizes plasmid stability can be found at intermediate values.

This observation indicates the existence of a non-linear relationship between stability of plasmids and the mean PCN of the population. To further explore this association, we computationally estimated the time-to-extinction in a long-term setting (simulations running up to 500 days) for different values of PCN and fitness cost. As expected, Figure 4C shows an accelerated rate of plasmid loss in costly plasmids. Crucially, there appears to be a critical PCN that maximizes the time-to-extinction, that depends on the per-cell plasmid cost. The time-to-extinction gives a notion of the stability of plasmids, but this measure may not apply if we introduce antibiotics, and therefore avoid plasmid extinction. For this reason, we also quantified plasmid stability by measuring the area under the curve (AUC) of simulation trajectories similar to those in Figure 4B. The heatmap illustrated in Figure 4D shows this measure highlighting the existence of a region in the cost-PCN plane, at intermediary PCN values, where plasmid stability is maximized.

### 2.2 Evaluating the role of selection in the stability of plasmids

To study the interaction between plasmid stability and the strength of selection in favor of PB cells, we assumed that the plasmid carries a gene that confers a selective advantage to the host in specific environments (e.g. resistance to heavy metals or antibiotics). For the purpose of this study, we will consider a bactericidal antibiotic (e.g. ampicillin) that kills PF cells with a probability that depends on the antibiotic dose. This results in a competitive advantage of the PB cells with respect to the PF subpopulation in this environment. We denote the intensity of this selective pressure by *α*.

Figures 5A-G illustrate plasmid dynamics over time for different values of *α*, obtained numerically by solving equation (2) with a fixed PCN (*n* = 19) and drug always present in the environment (*T* = 1). In our model, the we found a critical dose that stabilizes plasmids in the population, that is, the minimum selective *α, MSα* = *κ*_*n*_ + *μ*_*n*_(1 − *κ*_*n*_) (see Appendix B). The existence of a minimum selective concentration (MSC) that maintains plasmids in the population is a feature used routinely by bioengineers to stabilize plasmid vectors through selective media.^29^ Recall that the in our model the PF MIC is *α* = 1, therefore the *MSα* can be directly compared to the MSC/MIC ratio previously proposed^28, 30^ as a concern factor on selection of resistant strains in the environment.

**Figure 5.**
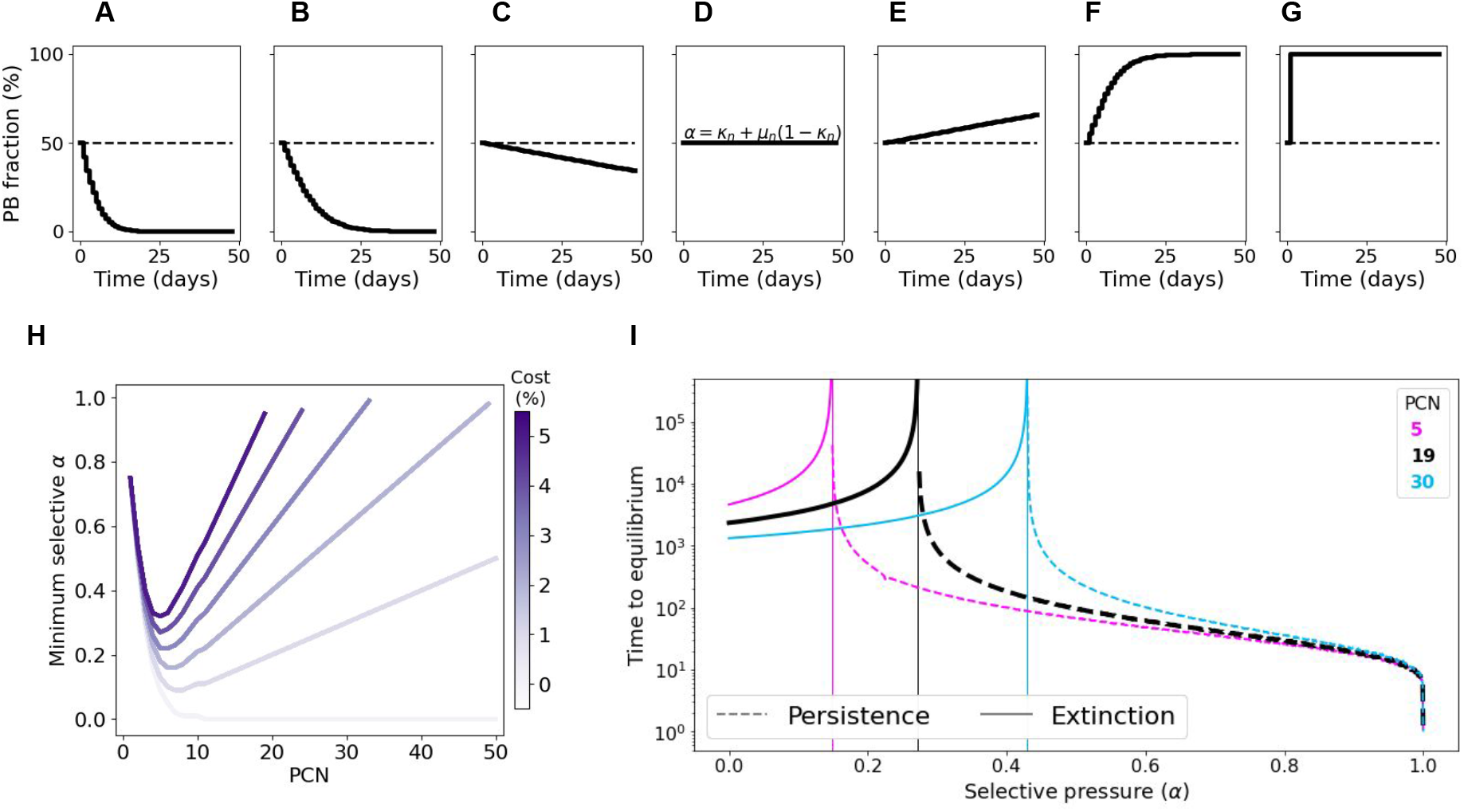
Numerical results illustrating the effect of a constant selective pressure in the stability of non-transmissible multicopy plasmids. **A-G)** Each box illustrates the temporal dynamics of the plasmid-bearing subpopulation in a pairwise competition experiment inoculated with equal initial fractions of *PF* and *PB*. From left to right, *α* = 0, 0.2, 0.26, 0.28, 0.6 and 1. The dotted line denotes *MSα* = *κ*_*n*_ + *μ*_*n*_(1 −*κ*_*n*_) for *n* = 19 and *κ*_*n*_ = 0.27. Note that for values of *α* < *MSα*, plasmids are unstable and eventually cleared from the population, while for *α* > *MSα* the plasmid-bearing subpopulation increases in frequency until reaching fixation. For *α* = *MSα*, the selective pressure in favor of the plasmid compensates its fitness cost and therefore the plasmid fraction remains constant throughout the experiment. **H)** Minimum selective pressure required to avoid plasmid loss for a range of PCNs. Different curves represent plasmids with different fitness costs (light purple denotes cost-free plasmids and dark purple a very costly plasmid). Note that, for costly plasmids, there exists a non-monotone relationship between *MSα* and PCN. **I)** Time elapsed before plasmid fraction in the population is stabilized, for different copy numbers (5 in magenta, 19 in black, and 30 in cyan). Dotted lines represent plasmid fixation, while dashed lines denote stable co-existence between plasmid-free and plasmid-bearing subpopulations, and solid lines plasmid extinction. The vertical line indicates *MSα*, the minimum selective pressure that stably maintains plasmids in the population. Black letters indicate the parameter values used in the examples shown in **A-G**.

As illustrated in Figure 5H, both low-copy and high-copy plasmids are inherently unstable and therefore the selective pressure necessary to stabilize them is relatively high, particularly for costly plasmids. Interestingly, at intermediate PCN values, the selective conditions necessary to stabilize plasmids are considerably less stringent than for low- and high-copy plasmids. This is the result of the non-linear relationship between MS*α* and *n*; since *μ*_*n*_ decreases exponentially with *n*, while *κ*_*n*_ increases only linearly with *n*.

Figure 5I shows the time elapsed before converging to a steady-state (either extinction or persistence) for different values of *α* and PCN. As *α* increases, the cost of plasmid-bearing is compensated by the benefit of carrying the plasmid and therefore plasmids are maintained in the population for longer. Note that at large values of *α*, plasmid-free cells are killed immediately independently of the mean PCN of the population, resulting very fast in a population composed almost exclusively of plasmid-bearing cells. Note that, in the case, the steady state 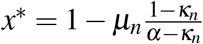 is achieved independently of the initial fraction of PB cells (see Appendix B), which is consistent with previous results^31^.

### 2.3 Plasmid stability in periodic environments

The purpose of this section is to understand the ecological dynamics of the plasmid-bearing population in fluctuating environments, i.e. when periodic antibiotic pulses are administered. We started by exploring the time duration a PB population can survive without antibiotics before being rescued by a strong antibiotic pulse (Figure 6A). Consistently with the results from the first section, lower plasmid costs result in increased rescue times, suggesting that a lesser rate of antibiotic exposure is required for their maintenance. In Figure 6B, we quantified this minimal period as a function of PCN and *α*. Note that higher values of *α* correspond to longer periods, which follows from the fact that a higher selective pressure increases the PB frequency. Figure 6D illustrates this critical period for *PCN* = 19.

**Figure 6.**
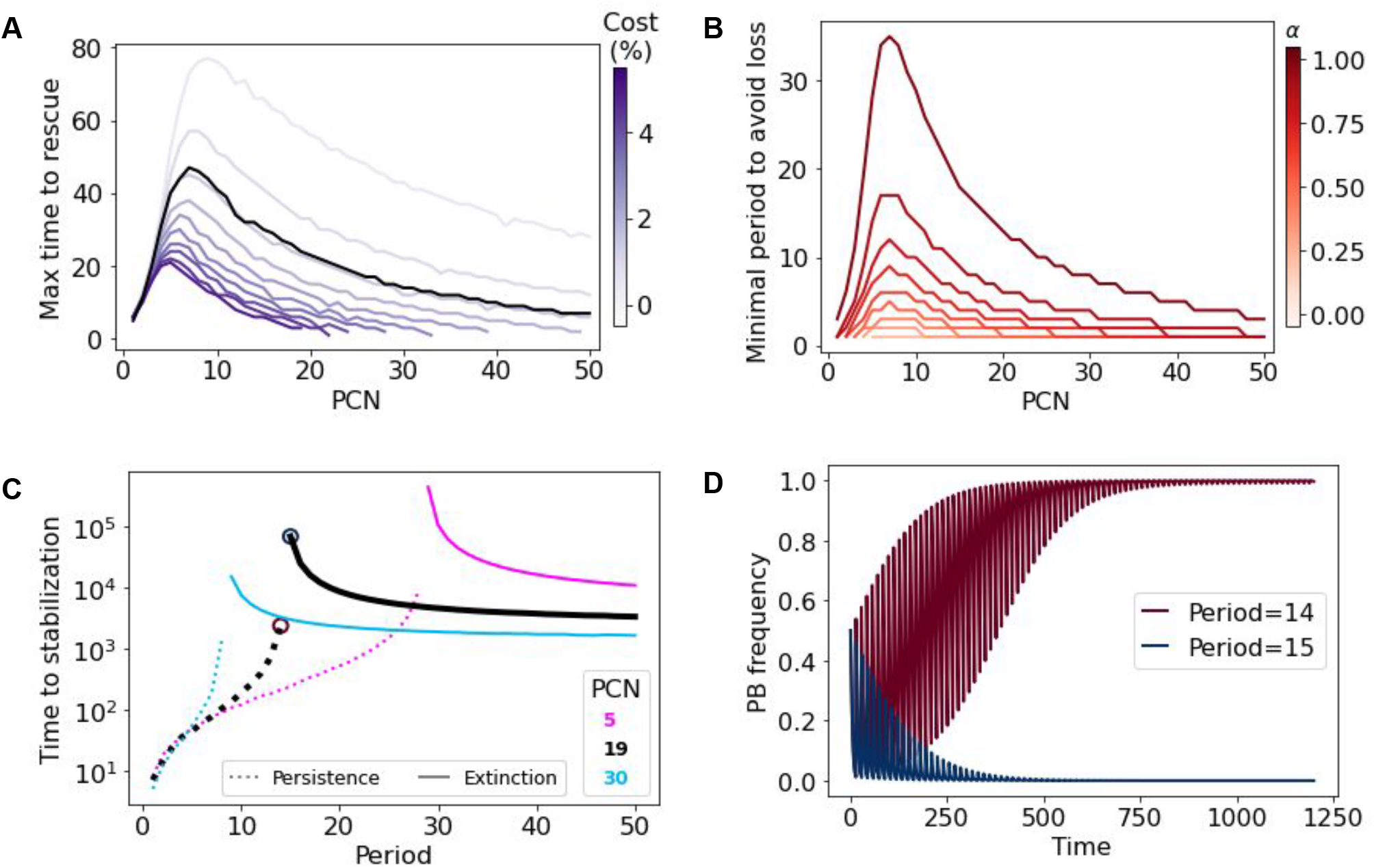
Numerical results of the model in periodic environments. **A)** Maximum time a plasmid population can grow without antibiotics to avoid plasmid loss when applying a strong antibiotic pulse. Curves represent how this time is affected by PCN. Blue intensity represents plasmid cost, and black line indicates results using the pBGT parameters. **B)** Minimal period required to avoid plasmid extinction. Simulations were performed using the pBGT measured cost (*κ* = 0.014). Red intensity represents different values of *α*. Note that higher values of *α* increase the minimal period. **C)** Time required for trajectories to stabilize for copy numbers 5, 19, and 30 using *α* = 0.99 and the measured cost per plasmid. Note that there is a critical period that defines fixation or coexistence marked by red and blue circles on the PCN=19 (black) curve. **D)** Trajectories for the critical periods of PCN=19 starting from 0.5 PB-PF frequency. Note that one day period difference leads to opposite outcomes.

In periodic environments, the relative abundance of the PB population is driven to zero (extinction) or reaches a steady state in which the plasmid fraction oscillates around an equilibrium frequency (persistence). In Figure 6C, times to stabilization were estimated for the strong selection regime (*α* = 0.99), using the same PCNs as in Figure 5I. Notice that the time-to-extinction is larger than the time to reach the periodic attractor. In both cases, the maximal time to rescue and the minimum period to avoid loss, we observe a non-monotone effect of PCN and, therefore, a range of PCNs whereby plasmid stability is maximized. This is consistent with what we observed without antibiotics (Figure 4C) and with constant environments (Figure 5H).

### 2.4 Optimal PCN depends on the rate of environmental fluctuation

In this section, we aim at exploring the concept of optimal PCN and how it depends on the environment. To do so, we define the optimal PCN (hereafter denoted PCN^*^) as the PCN that maximizes the area under the curve (AUC) of the PB frequency over time. This notion of stability was already introduced in Figure 4D and has the advantage that it can be used when the PB fraction goes to 0, to a fixed equilibrium, or when it oscillates.

First, we calculated PCN^*^ for a range of plasmids fitness costs in the absence of selection (black solid line of Figure 7A) and found that PCN^*^ is inversely correlated with the plasmid fitness cost. In order to compare the optimal PCN predicted by the model with PCN values found in other experimental plasmid-host associations, we searched the literature for studies that measure both PCN and fitness cost. These values are summarized in Table 3 and illustrated in Figure 7A. The values of PCN found in the literature were below the predicted PCN^*^ in an antibiotic-free regime (black solid line), suggesting that plasmids would be unstable in the absence of selection. But, crucially, PCN values obtained from the literature are within the blue-shaded area that represents the PCN^*^ estimated for different environments (observe the non-linear relationship between *α*, PCN^*^, and cost, in line with our previous findings).

**Table 3.** Plasmid copy number and plasmid costs from literature. SD Standard Deviation.

| Name | Plasmid Type | Species | PCN | PCN SD | Cost | Cost SD | Reference |
| --- | --- | --- | --- | --- | --- | --- | --- |
| pB1006 | ColE1 | *Haemophilus influenzae* RdKW20 | 10.530 | 1.112 | 0.021 | 0.012 | 58 |
| pB1005 | ColE1 | *Haemophilus influenzae* RdKW20 | 20.450 | 2.590 | 0.05 | 0.013 | 58 |
| pB1000 | ColE1 | *Haemophilus influenzae* RdKW20 | 25.020 | 1.920 | 0.054 | 0.002 | 58 |
| pNI105 |  | *Pseudomonas aeruginosa* | 18.000 | 2.400 | 0.132 | 0.025 | 59 |
| 2-uM |  | *Saccharomyces cerevisiae* | 52.000 | 0.000 | 0.0884 | 0.00416 | 17 |
| pNUK73 |  | *Pseudomonas aeruginosa* | 11.030 | 1.890 | 0.214 | 0.008 | 59 |
| pBGT | ColE1 | *Escherichia coli* | 19.120 | 1.560 | 0.057 | 0.013 | 22 |
| pBGT R164S | ColE1 | *Escherichia coli* | 21.100 | 0.850 | 0.057 | 0.003 | 22 |
| pBGT G54U | ColE1 | *Escherichia coli* | 44.500 | 3.810 | 0.207 | 0.019 | 22 |
| pBGT G55U | ColE1 | *Escherichia coli* | 88.930 | 15.650 | 0.443 | 0.116 | 22 |
| pBGT R164S G54U | ColE1 | *Escherichia coli* | 52.300 | 2.190 | 0.238 | 0.016 | 22 |
| pBGT R164S G55U | ColE1 | *Escherichia coli* | 127.290 | 4.580 | 0.491 | 0.082 | 22 |

**Figure 7.**
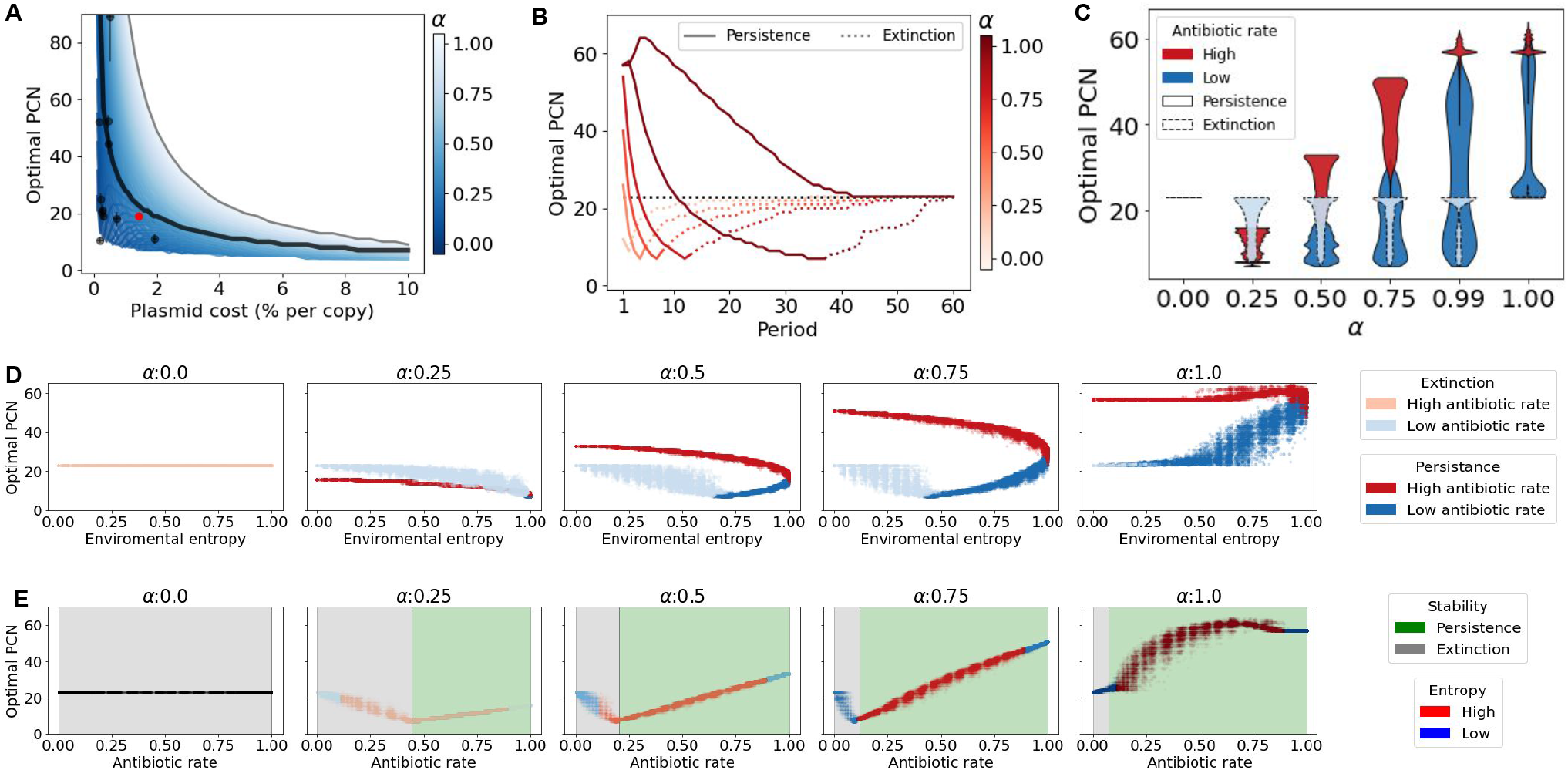
Optimal PCNs in fluctuating environments. **A)** Optimal plasmid copy number (PCN^*^) as the number of copies that maximizes the area under the curve of Figure 4B. PCN^*^ decreases exponentially as we increase the fitness cost associated with carrying plasmids, as indicated in black-solid line. Black dots show some PCN-costs data obtained from the literature. Red dots indicate the values of pBGT. Blue-scale lines indicate optimal PCN curves for many values of *α*. Light-blues indicate higher values of *α* whereas dark-blues indicate lower values of *α*. Gray line shows the max PCN for the corresponding plasmid cost. **B)** Optimal PCN in periodic environments. Each curve corresponds to a value of *α*. Black line shows *α* = 0. Observe that for very short periods optimal PCNs are high, then for certain period the optimal PCN reaches a minimum then as period increases, the optimal PCN tends to the optimal of *α* = 0. **C-E)** Optimal PCNs using random environments. **C)** Environments are classified by their rate of days with antibiotics, the rate differences produce a multi-modal outcome, where higher rates increases the optimal PCN and vice-versa. Simulations using the same environments were made for different *α*s. Note that *α* intensity increases the separation of the modes. Modes are also classified by their stability, persistence marked with a solid border line and extinction with a dashed border line. **D)** Panel of optimal PCNs plotted by the environment entropy for sample *α*. Environments are classified by their antibiotic rate. **E)** Panel of optimal PCNs plotted by the environment antibiotic rate for sample *α*. Environments are classified by their entropy.

These observations would be consistent with the constant use of antibiotics at low doses that reduces the optimal PCN. However, similar PCN^*^ values can be achieved by administering higher doses of antibiotics periodically, as illustrated in Figure 7B for the case of pBGT. Notice again the non-linear relationship between PCN^*^ and the frequency of antibiotic exposure. At very low frequencies, the PB population goes extinct before the first antibiotic pulse and intermediate PCNs maximize the AUC as in Figure 4D. At high antibiotic frequencies, the PB population persists and oscillates around some value that increases with PCN. This is consistent with a previous experimental study that evaluated the stability of costly plasmids in terms of the frequency of environmental fluctuation.^32^

Periodic environments provided us with insights into how selection acts on the mean PCN of the population, but natural environments are not periodic but randomly alternate between intervals of positive and negative selection. The role of environmental stochasticity in the stability of multicopy plasmids^23, 33^ and, in general, in the population dynamics of asexual populations has been widely studied.^34, 35^ In our model, we generated stochastic environments that randomly switch from antibiotic-free to antibiotic for a period of 1, 000 days. Each random environment is represented by a sequence of 1s and 0s, corresponding to days with and without antibiotics, respectively. Therefore stochastic environments can be characterized by their Shannon’s entropy (environmental entropy, H) and the fraction of days with drug exposure (antibiotic rate, AR) (see Appendix C). Environments were classified into “High” and “Low” depending on whether the AR was greater or lower than 0.5. Mind that each value of H corresponds to two AR values *AR* and 1 − *AR*.

Panels on Figures 7D-E show the PCN^*^ found by applying the stochastic environments ordered by entropy (or by AR), for different values of *α*. For low values of *α*, only high antibiotic rates lead to plasmid persistence. Notice the non-linear relationship between PCN^*^ and AR, similar to the observed for the period in the deterministic setting; PCN^*^ decreases with AR at low values (corresponding to extinction) but increases with AR at high values (corresponding to persistence). For higher values of *α*, we observed that high AR always leads to persistence, while low AR can lead to extinction if entropy is low. In fact, these low values of the entropy corresponded to long periods without antibiotics that drove the PB population to extinction. Another interesting remark is that the distribution of obtained PCN^*^s is multi-modal; at fixed entropy, plasmid persistence is achieved by high values of AR that correspond to high PCN^*^, or by low values of AR that correspond to a small value of PCN^*^. Similarly, a fixed value of *α* corresponds to two values of PCN^*^ depending on the antibiotic rate (Figure 7C).

## 3 Discussion

In this work, we used a population genetics modeling approach to study how non-transmissible plasmids are maintained in bacterial populations exposed to different selection regimes. In particular, we considered a small multicopy plasmid that lacks an active partitioning mechanism and therefore segregates randomly upon cell division. Multicopy plasmids are prevalent in clinical bacteria and usually carry antimicrobial resistance genes that can be transferred between neighboring bacterial cells,^36^ as well as other evolutionary benefits that go well beyond horizontal transfer.^37^ For instance, as multicopy plasmids are present in numerous copies per cell, the mutational supply increases proportionately and, once a beneficial mutation appears, its frequency can be amplified during plasmid replication. This results in an accelerated rate of adaptation to adverse environmental conditions^7^ and enables evolutionary rescue.^38^ Also, multicopy plasmids increase the genetic diversity of the population, thus enhancing survival in fluctuating environments^25^ and allowing bacterial populations to circumvent evolutionary trade-offs.^23^

While the benefits of carrying plasmids may be clear under certain circumstances, their maintenance can be associated with a considerable energetic cost in the absence of selection for plasmid-encoded genes. This trade-off between segregational stability and fitness cost has been shown to drive ecological and evolutionary dynamics in plasmid-bearing populations,^39^ resulting from multi-level selection acting on extrachromosomal genetic elements.^40, 41^ Plasmid population dynamics resulting from random segregation and replication result in a complex interaction between plasmid copy number, genetic dominance, and segregational drift, with important consequences in the fixation probability of beneficial mutations^42^ and the repertoire of genes that can be carried in mobile genetic elements.^43^ Besides a reduction in segregational instability, increasing the number of plasmids each cell carries also results in an increase in gene dosage^44, 45^ and expression variability of plasmid-encoded genes.^25, 46^ For this reason, plasmid control in wild-type bacteria is a tightly regulated process^47^ that depends on the environment and the host’s genetics.^8^ Precise PCN control is also an important feature of synthetic genetic circuits that use plasmids as vectors for the production of recombinant substances.^48^

To explore the interaction between the strength of selection and PCN, in this manuscript we postulated discrete-time and Wright-Fisher diffusion models with the following biological assumptions:1) Plasmids encode for accessory genes that confer an advantage in harsh environments, for instance, antibiotic resistance genes; 2) Bearing plasmids is associated with a fitness cost in the absence of selection for plasmid-encoded genes; 3) Each plasmid segregates randomly to a daughter cell upon division; thus, plasmid bearing bacteria can produce plasmid-free cells with a probability of 1*/*2^*n*^, where *n* is the PCN; 4) The cost associated with plasmid-bearing is constant in time (no compensatory adaptation). We parameterized the model using a well-characterized multicopy plasmid, pBGT,^22–24^ and estimated the maximal growth rates of plasmid-bearing and plasmid-free cells by analyzing growth kinetics of each strain grown in isolation. From the growth curves, we obtained estimates for the fitness cost associated with plasmid bearing and the fitness advantage of the plasmid-bearing cells for a range of antibiotic concentrations. We also performed one-day competition experiments between different subpopulations of PB and PF cells and evaluated how this fraction changed after a day of growth in media supplemented with antibiotics. Using this approach we obtained theoretical and experimental iterative maps that we used to predict the long-term dynamics of the system.

Altogether, our results suggest that plasmid population dynamics in bacterial populations is predom-inantly driven by the existence of a trade-off between segregational loss and plasmid cost. We found that selection is necessary for the persistence of costly plasmids in the long term, and that the strength of selection is highly correlated with the final fraction of plasmids in the entire population. As a result, whether plasmids are maintained or lost in the long term results from the complex interplay between PCN and its fitness cost, as well as the intensity and frequency of positive selection. As shown in the exhaustive exploration of parameters performed in this study, these relationships are highly non-linear, thus resulting in the existence of an optimal PCN that depends on the rate of environmental fluctuation, the number of plasmids carried in each cell, and the fitness burden conferred by each plasmid-encoded gene in the absence of selection. In random environments, we observed a bimodal PCN^*^ distribution, similar to the plasmid size distribution described for non-transmissible plasmids^49^ and for conjugative plasmids.^50^

Although both our theoretical and experimental models consider a multicopy plasmid with random segregation, the existence of an optimal PCN should also hold for non-random segregation (e.g. active partitioning), as this would decrease the probability of segregational loss (which corresponds to having a smaller value of *μ*_*n*_ in our model) so its optimal copy number will likely be lower than a plasmid that relies on random segregation.^51^ In contrast, compensatory adaptation that reduces the fitness cost associated with plasmid bearing (in our model, a lower value of *κ*_*n*_), would result in an increase in PCN^*^. We conclude by arguing that, as the existence of plasmids in natural environments requires intermittent periods of positive selection, the presence of plasmids contains information on the environment in which a population has evolved. Indeed, the plasmid copy number associates the frequency of selection with the energetic costs of plasmid maintenance. That is, there is a minimum frequency of drug exposure that allows multiple copies to persist in the population, and, for each environmental regime, there is an optimal number of plasmid copies.

## 4. Appendix A: Experimental methods

### 4.1 Bacterial strains and media

The plasmid free strain we used was *E. coli* K12 MG1655 and the plasmid bearing strain was MG/pBGT carrying the multicopy plasmid pBGT with the *β* -lactamase *bla*_TEM-1_ which confers resistance to ampicillin and the fluorescent protein GFP under an arabinose inducible promoter. Mean plasmid copy number in the population is *PCN* = 19.1 ± 3.8.^22^ Overnight cultures were grown in flasks with 20 ml of lysogeny broth (LB) (Sigma L3022) with 0.5 % w/v L-(+)-Arabinose (Sigma A91906) for fluorescence induction, in a shaker-incubator at 220 RPM at 37 ^°^C. For the plasmid bearing strain, 25 mg/l of ampicillin (Sigma A0166) were added to eliminate segregant cells. Ampicillin stock solutions were prepared at 100 mg/ml directly in LB and sterilized by 0.22 *μ*m (Millex-GS SLGS033SB) filtering. Arabinose stock solutions were prepared at 20% w/v in DD water and sterilized by filtration.

#### 4.2 Bacterial growth experiments

Growth kinetics measurements of each strain were performed in 96 well plates with 200 *μ*l of LB with 0.5% w/v arabinose without antibiotics, plates were sealed using X-Pierce film (Sigma Z722529), each well seal film was pierced in the middle with a sterile needle to avoid condensation. Plates were grown at 37 deg*C* and reading for OD and fluorescence were made every 20 minutes in a fluorescence microplate reader (BioTek Synergy H1), after 30 seconds linear shaking.

#### 4.3 Competition experiments

Competition experiments were performed using 96-well plates with 200 *μ*l of LB with 0.5% w/v arabinose, and respective ampicillin concentrations: 0, 1, 2, 2.5, 3, 3.5, 4, and 6 mg/l was implemented by plate rows. To construct our inoculation plate, overnight cultures of the plasmid-free strain and the plasmid bearing strain were adjusted to an OD of 1 (630 nm) using a BioTek ELx808 Absorbance Microplate Reader diluted with fresh ice cooled LB. Appropriate volumes were mixed to make co-cultures at fractions 0, 0.1, 0.2, …, 1 and set column-wise on a 96-well plate (Corning CLS3370). We then used a 96 pin microplate replicator (Boekel 140500) with flame sterilization before each inoculation. Four replicates plates were grown in static incubator at 37^°^*C*. After 24 hours growth, plates were read in a fluorescence microplate reader (BioTek Synergy H1) using OD (630 nm) and eGFP (479,520 nm) after 1 minute of linear shaking.

#### 4.4 Plasmid fraction determination

To calculate the fluorescence intensity, we first subtracted the background signal of LB for fluorescence and OD respectively, then the debackgrounded the fluorescence signal was scaled by dividing by the debackgrounded OD. The measurements for our inoculation plate showed a strong linear correlation (*R*^2^ = 0.995) between co-cultures fractions and fluorescence intensity (Figure 3B). This allowed to directly approximate the populations plasmid fractions from the readings of our competition experiments. We normalized the data independently for each antibiotic concentration taking the average measurements of the 4 replicates. Plasmid fractions, *PF*, were inferred by normalizing the mean fluorescence intensity for each well, *f*_*i*_, to the interval [0,1] using the following formula: *PF*_*i*_ = (*f*_*i*_ − *f*_*min*_)*/*(*f*_*max*_ − *f*_*min*_) were *f*_*max*_ and *f*_*min*_ are the mean fluorescence intensities at fractions 1 and 0 respectively.

## 5 Appendix B: Mathematical model

### 5.1 Fixed points of equation (2)

Let 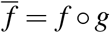. We want to study the fixed points of 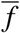 and their domains of attraction. It is not hard to see that 0 is always fixed point, and once the frequency reaches 0 it stays at 0. In addition, if *x* ≠ 0,

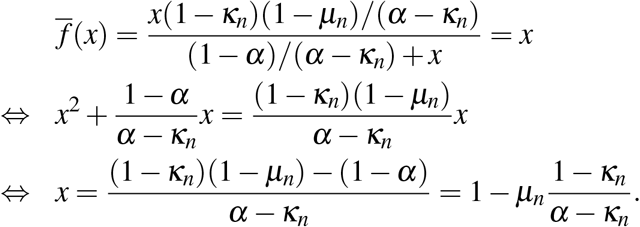

Denote *x*^*^ := 1 − *μ*_*n*_(1 − *κ*_*n*_)*/*(*α* − *κ*_*n*_). Since the frequencies are in [0, 1], this fixed point only exists if *α > κ*_*n*_ + *μ*_*n*_(1 − *κ*_*n*_). As *n* increases, *μ*_*n*_ decreases exponentially, while *κ*_*n*_ increases only linearly, so there is a non linear relationship between *n* and the minimum *α* required for the existence of a second fixed point *x*^*^.

Let us analyze the stability of *x*^*^. Let us assume that *α > κ*_*n*_ + *μ*_*n*_(1 − *κ*_*n*_).

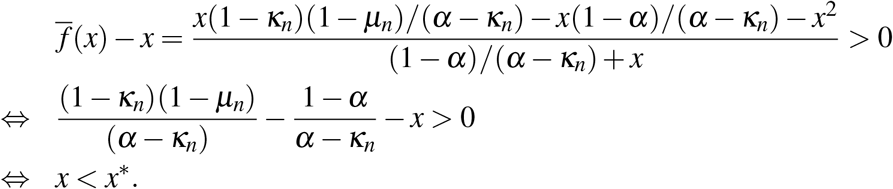

So, the frequency increases if it is below *x*^*^ and decreases otherwise, meaning that it is a stable fixed point. In addition, the domain of attraction is (0, 1], meaning that this equilibrium fraction is reached for any initial state.

To sum up, 0 is always a fixed point. If *α > κ*_*n*_ + *μ*_*n*_(1 − *κ*_*n*_) then there is an additional stable fixed points *x*^*^.

#### 5.2 Choice of the model

In this section, we compare two types of mathematical models for the evolution of plasmid-bearing frequencies, the discrete time model used in this paper (eq (2)) and the Wright-Fisher diffusion.

There had been several attempts to adapt the classical theory of Wright Fisher models to this experimental setting (see for example^52^). A mathematical rigorous way to do this was developed in.^19^ In^53^ an heuristic and applicable to data framework was introduced. Recently, in,^54^ the two methodologies had been paired in order to have a rigorous and applicable way to use classic population genetics to study evolutionary experiments. In this work, days take the role of generations, and as the number of individuals after each sampling is more or less constant, the assumption of constant population size becomes reasonable.

Let us assume that the mutation rate *μ*_*N,n*_ = 2^−*n*^ and the cost *κ*_*N,n*_ are parameterized by *N*. To see the accumulated effects of plasmid costs, segregational loss and genetic drift, we need *κ*_*N,n*_ and *μ*_*N,n*_ to be of order 1*/N* (see e.g. Chapter 5 in^55^). The first condition is fulfilled if the cost per plasmid is very low, for example when *κ*_*N,n*_ = *κn/N*. The second one stands if *n* is of order log_2_(*N*), which is the case, for example, if *n* = 20 and *N* = 10^6^, or if *n* = 15 and *N* = 10^5^. In that case we set *μ* = *N*2^−*n*^. Under this setting, when time is accelerated by *N*, the frequency process of individuals with plasmids can be approximated by the solution of the stochastic differential equation (SDE)

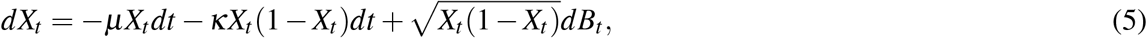

where *B* is a standard Brownian motion. This is known as the Wright-Fisher diffusion with mutation and selection. When antibiotic is added, at times {*T*, 2*T*,… }, then (5) modifies to

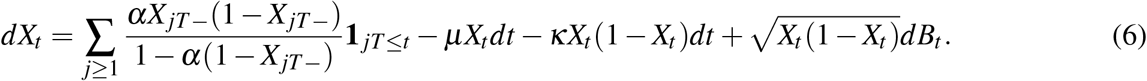

However, in our experimental setting, the cost that we measure (*κ*_*n*_ ≃ 0.27) is much higher than the inverse population size, so we are in the regime of strong selection. In other words, for plasmids that have a very small cost, of the order of 1*/N*, genetic drift would play an important role, and the above Wright-Fisher diffusion with mutation, selection and antibiotic peaks (6) would be the most suitable model. But in our setting, selection (plasmid costs) is so high that genetic drift becomes negligible. Recall that equation (2) does not need any time rescaling, whereas, in the diffusion (6) time is measured in units of *N* generations. Under strong selection, the frequencies evolve much faster.

## 6 Appendix C: Numerical simulations

### 6.1 Computer implementation

The model was implemented in Python, using standard scientific computing libraries (Numpy, Mat-plotLib, and the Decimal library was required to resolve small numbers conflicts). In general, all simulations started at PB frequency 1 (unless stated otherwise). Numeric simulations were defined to reach a steady state when values first repeat. In the case of periodic environments, the repetition must happen at antibiotic peaks days. We considered extinction if the end point of the realization dropped below a threshold adjusted to the simulations times, the highest being 1 × 10^−7^ and the lowest 1 × 10^−100^.

#### 6.2 Random environments

Environmental sequences of size 1000 (days) using a binomial distribution varying the probability of success. For each environment created we also bit-flipped (so 101… turns into 010…) and two measures was applied to each resulting environments. First, we used Shannon entropy, 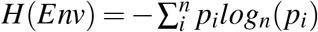, with two states, *n* = 2 (antibiotic or no-antibiotic) and *p*_*i*_ equal to the probability of finding an state day, i.e. the fractions of days with antibiotics and without antibiotics. We classified environments by their *H* and by the fraction of antibiotic days, as being this an important feature. This two measures are in the [0,1] interval so we binned the intervals into 20 bins and 1,000 environments were created for each bin.

#### 6.3 Model parametrization

Growth kinetics parameters were estimated using the R^56^ package growth rates.^57^ Exponential phase duration, *σ*, was calculated by finding lag phase duration and the time to reach carrying capacity using the non-linear growth model Baranyi. Maximum growth rates, *r* and *r* + *ρ*_*n*_, were estimated using the Nonparametric smoothing splines method. *κ*_*n*_ value was estimated using equation (1) and the data from the antibiotic-free competition experiment using a curve fitting algorithm from the SciPy library in a custom Python script. Respective values of *α* were found in the same manner using equation (2) and fixing *κ*_*n*_. *κ*_*n*_ was also calculated using the formula in equation 4 with a very similar result. The parameters are summarized in Tables 1 and 2.

